# NMR reveals light-induced changes in the dynamics of a photoswitchable fluorescent protein

**DOI:** 10.1101/692087

**Authors:** N. E. Christou, I. Ayala, K. Giandoreggio-Barranco, M. Byrdin, V. Adam, D. Bourgeois, B. Brutscher

## Abstract

The availability of fluorescent proteins with distinct phototransformation properties is crucial for a wide range of applications in advanced fluorescence microscopy and biotechnology. To rationally design new variants optimized for specific applications, a detailed understanding of the mechanistic features underlying phototransformation is essential. At present, little is known about the conformational dynamics of fluorescent proteins at physiological temperature, and how these dynamics contribute to the observed phototransformation properties. Here, we apply high-resolution NMR spectroscopy in solution combined with in-situ sample illumination at different wavelengths to investigate the conformational dynamics of rsFolder, a GFP-derived protein that can be reversibly switched between a green fluorescent state and a non-fluorescent state. Our results add a dynamic view to the static structures obtained by X-ray crystallography. Including NMR into the analytical toolbox used for fluorescent protein research provides new opportunities for investigating the effect of mutations or changes in the environmental conditions on the conformational dynamics of phototransformable fluorescent proteins, and their correlation with the observed photochemical and photophysical properties.

**Significance:** Photo-transformable Fluorescent Proteins (PTFPs) are essential tools for super-resolution (SR) microscopy. In practical applications, however, researchers often encounter problems when using PTFPs in a particular cellular context, because the environmental conditions (pH, temperature, redox potential, oxygen level, viscosity, …) affect their brightness, photostability, phototransformation kinetics, etc. Rational fluorescent protein engineering exploits the mechanistic information available from structural studies, mainly X-ray crystallography, in order to design new PTFP variants with improved properties for particular applications. Here we apply NMR spectroscopy in solution to investigate the light-induced changes in conformational dynamics of rsFolder, a reversibly switchable fluorescent protein. The dynamic view offered by NMR highlights protein regions that comprise potentially interesting mutation points for future mutagenesis campaigns.

## Introduction

The discovery and engineering of photo-transformable fluorescent proteins (PTFPs), derived either from *Aequora victoria* green fluorescent protein (GFP) or other fluorescent proteins from Anthozoa species (corals and anemones), has triggered exciting new modalities in fluorescence microscopy (1) and biotechnology (2). In particular, PTFPs have become indispensable tools for modern super-resolution light microscopy techniques that achieve resolution beyond the diffraction limit (3, 4). At present, a large number of PTFPs have been discovered or engineered that differ in their specific response to light (photoactivation, photoconversion, or reversible photoswitching), in the kinetics of this response, in their photostability and general fluorescence characteristics (brightness, emission color, Stokes shift, …), as well as in their oligomerization state, folding stability, and maturation efficiency. Furthermore, these properties are generally dependent on environmental conditions (5). The requirements for an optimal PTFP marker are different for specific applications, which calls for further expansion and improvements of the PTFP toolbox.

Understanding at near-atomic resolution the mechanistic features responsible for the observed photochemical and photophysical properties of PTFPs is a key step to envision the rational design of new variants optimized for a particular application or experimental context. Most mechanistic studies on PTFPs reported in the literature have used X-ray crystallography as the method of choice to determine structures of their different ground-state conformations. Such structures are essentially static and, as data are typically collected at cryogenic temperature, they are representative of frozen states. Kinetic crystallography has also been used in combination with optical spectroscopy and/or molecular dynamics simulations to decipher the structures of short-lived ground state species such as intermittent dark states (6, 7). Recently, ultrafast time-resolved crystallography using an XFEL source has been able to image an electronically excited short-lived intermediate state in rsEGFP2 that becomes populated during photoswitching between a nonfluorescent (“off”) and a fluorescent (“on”) state (8), shedding some light on the switching dynamics of a PTFP at the picosecond timescale.

While FPs are generally quite resistant to their environment, an increasing number of studies have shown that PTFP photophysics are affected by external factors such as pH, redox potential, oxygen level, viscosity, temperature, etc (5, 9–13). These environmental effects are difficult to study by X-ray crystallography where structural data are collected in conditions imposed by crystallogenesis that may differ substantially from the conditions found during in-situ fluorescence imaging. Moreover, little is known about the conformational dynamics of PTFPs at physiological temperatures, and their relevance for the observed fluorescence and phototransformation properties. To address these issues, multidimensional liquid-state NMR spectroscopy may provide a complementary tool to investigate at atomic resolution the conformational dynamics of PTFPs in different states, and over a wide range of temperatures and buffer conditions. However, quite surprisingly, so far only little information on PTFPs has been obtained from NMR (11, 14, 15), while a few more studies on chromophore dynamics in conventional GFPs can be found in the literature (16–18).

Here, we introduce a toolbox of multidimensional and multinuclear (^1^H, ^13^C, ^15^N) NMR experiments specifically tailored to investigate PTFPs structural dynamics. We apply this toolbox to probe the conformational dynamics of rsFolder, a PTFP that can be reversibly switched from a stable fluorescent “on” state to a meta-stable nonfluorescent “off” state, and vice versa, upon illumination with cyan or violet light, respectively. Such reversibly photo-switchable fluorescent proteins (RSFP) are required for a variety of applications, ranging from super-resolution microscopy (RESOLFT, NL-SIM, pcSOFI, and others) (19–21), optical lock-in detection (OLID) (22), multiplexed imaging (OPIOM) (23), up to the development of high-density optical storage media (2). rsFolder has been genetically engineered as a chimera between rsEGFP2 (24) and superfolder GFP (25) to facilitate proper folding in highly reducing environments such as the bacterial periplasm (26). The photophysical properties of rsFolder have been characterized by optical spectroscopy, and high resolution crystallographic structures are available for both the “on” and “off” states (26). These structures reveal the typical 11-stranded β-barrel fold, enclosing an endogenous *p*-HBI chromophore, formed by maturation of an Ala-Tyr-Gly tripeptide. Upon light illumination at 488 nm (405 nm), the chromophore undergoes a *cis* to *trans* (*trans* to *cis*) isomerization, and protonation (deprotonation) at the ζ position of the phenol ring. While the β-barrel structures for the “on” and “off” states almost perfectly superpose (C^α^ rmsd = 0.21 Å), slight structural rearrangements are observed for a few side chains located in the close vicinity of the chromophore (Figure 1). Our NMR data complement this static picture by adding information on the changes in local conformational sampling after photoswitching, providing some new insight toward the long-term goal of better understanding how the photophysical properties of PTFPs are influenced by different types of local molecular flexibility. This work opens the door to further studies aiming at evaluating how PTFPs conformational dynamics respond to mutational or environmental changes.

**Figure 1.**
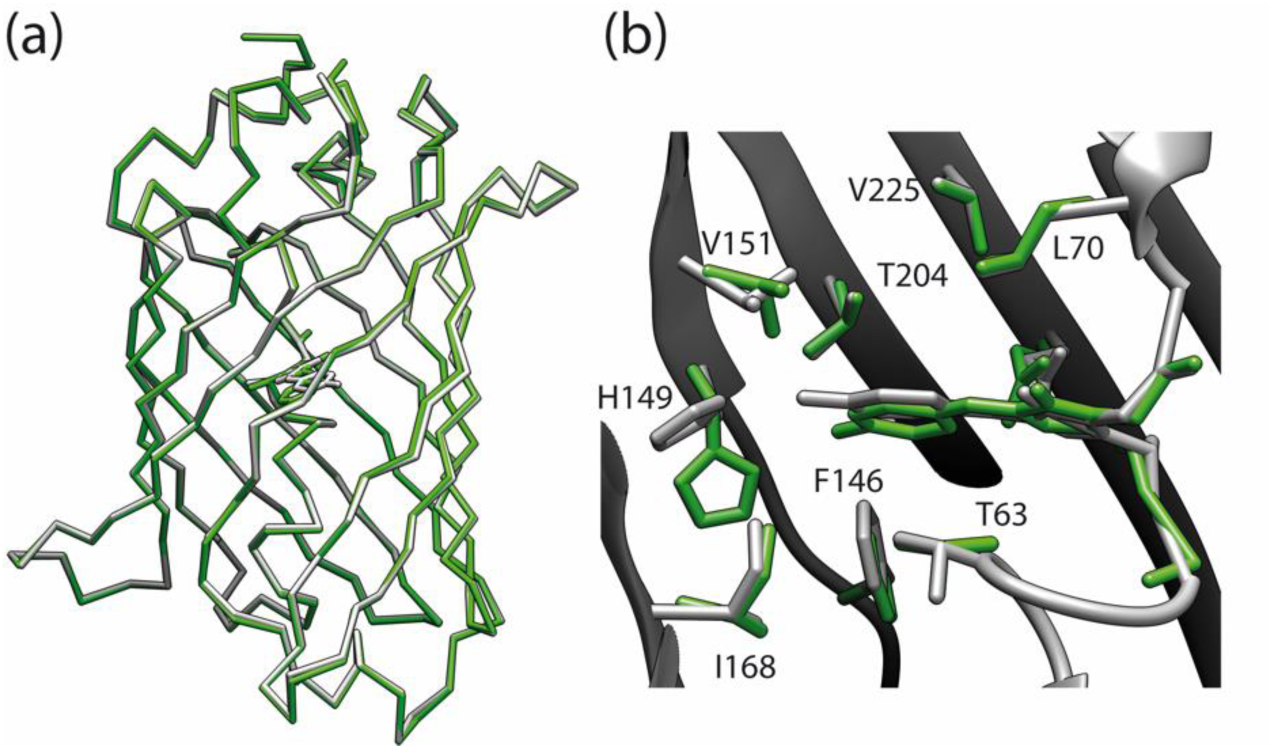
Crystallographic structures of rsFolder in the fluorescent “on” state (green) and the non-fluorescent “off” state (grey). (a) Superposition of the main chain coordinates highlighting that the overall β-barrel structure of the protein is identical in the two states. (b) Zoom on the chromophore region of rsFolder showing the change in chromophore geometry from “cis” to “trans”, as well as some side chain rearrangements in the vicinity of the chromophore. The most pronounced structural changes upon light illumination are observed for F146, H149 and V151.

## Materials and Methods

### Protein samples

Purification of rsFolder was performed as described previously (26). The full-length rsFolder cDNA was cloned in the ampicillin-resistant expression plasmid pET15b (Addgene, Teddington, UK), and constructs were designed to bear a six-residue N-terminal His-tag linked to the protein coding region via a Thrombin cleavage sequence. When considered necessary, the His-tag was cleaved using a Thrombin protease. *Escherichia coli* BL21(DE3) cells were transformed with the plasmid described above for protein expression. An isotopically enriched minimal medium, containing ^15^NH_4_Cl (1 g/L) and ^13^ C-glucose (2 g/L), was used to grow the cells until the cultures reached an OD at 600 nm between 0.6 and 0.8, at which point protein expression was induced by adding IPTG (isopropyl β-D-1-thiogalactopyranoside). For uniformely ^15^N/^13^C labeled samples, 1mM IPTG was added and induction was performed overnight at 20°C. The cells were harvested by centrifugation and subsequently lysed by sonication in lysis buffer (50 mM HEPES, 100 mM NaCl, pH 7.5). The lysate was centrifuged at 4°C for 30 min at 46 000g. The protein present in the supernatant was purified using a Ni-NTA column (Qiagen). The protein was eluted by increasing the concentration of imidazole. The resulting protein was further purified using size exclusion chromatography with a Superdex 75 column (GE) equilibrated with the NMR buffer (50 mM HEPES pH 7.5).

### In-situ laser illumination inside the NMR magnet

An in-situ laser illumination device similar to that described by Mizuno et al. (15) was built. It comprises an Oxxius laser box (wavelength combiner L4Cc, Oxxius S.A., Lannion, France) hosting two lasers of wavelengths 405 nm (LBX-405-180-CSB-OE) and 488 nm (LBX-488-200-CSB-OE), and an optical fiber (Thorlabs FG105UCA, 0.22NA, 105 μm core diameter, 2 mm diameter protective cover, 5 m length). The fiber end was left naked and routinely cleaved when necessary, using a Ceramic Fiber Scribe (Thorlabs CSW12-5). The fiber was inserted into the plunger of a 5 mm diameter Shigemi tube (Shigemi Co LTD, Tokyo, Japan). Importantly the Shigemi plunger was filled with water before inserting the fiber to avoid a sudden change in the refractive index at the air-glass interface, thus reducing the divergence of the laser beam. The laser power was set and monitored using the Oxxius Software from a laptop connected to the laser combiner, and the laser output was controlled either manually using a shutter, or by digital on/off modulation from the NMR console connected to the laser box via TTL lines. The difference between nominal laser power as displayed by the Oxxius software and the measured output power at the fiber tip was about 30%, suggesting a coupling efficiency of about 70%. For short illumination times (less than 1 minute), the full laser power was used (200 mW nominal), while for longer exposure times (>> 1 minute) the output was set to 30% of the maximal power to limit the effect of photobleaching. In order to allow smooth insertion of the fiber-coupled Shigemi tube into the NMR magnet, the use of 2 Bruker sample holders put on top of each other proved to be helpful.

### NMR measurements

All NMR experiments were performed on Bruker Avance IIIHD spectrometers, at the high-field NMR facility of IBS, operating at magnetic field strengths of 600 MHz (H/D exchange), 700-MHz (^15^N relaxation, NMR assignment), and 850-MHz (LD-EXSY, ^15^N CPMG, H/D exchange, methyl side chain, aromatics and chromophore NMR experiments), and equipped with cryogenically cooled triple-resonance probes and pulsed z-field gradients. NMR measurements were performed at 40°C on NMR samples containing 100 - 200 μM [U-^13^C, ^15^N]-enriched rsFolder in 50 mM HEPES (pH 7.5), and 5% (v/v) D_2_O. The samples were stored at 4 °C between data recordings, and the sample quality was routinely checked for possible aggregation and/or degradation by recording ^1^H-^15^N correlation spectra. “On” state data were recorded after short (5 s) illumination at 405 nm in the absence of laser illumination, while ‘off” state data were acquired after a short (10 s) high-power illumination at 488 nm, under continuous illumination (488 nm) using reduced (30%) laser power to prevent on-state recovery by thermal relaxation. Most NMR experiments used in this study are implemented in the NMRlib pulse sequence library for Bruker spectrometers (27) that can be freely downloaded from our web site (http://www.ibs.fr/research/scientific-output/software/pulse-sequence-tools). NMR data processing was performed using Topspin 3.5 (Bruker BioSpin), while data analysis was done using the CCPNMR software (28).

For sequential resonance assignment of the “on” state, a set of five triple-resonance 3D BEST-TROSY experiments (29, 30), HNCO, HNcoCA, HNCA, HNcoCACB, and HNCACB, was recorded. For the “off” state, two LD-EXSY data sets (^15^N,^15^N,^1^H) and (^1^H,^15^N,^1^H) were recorded to transfer the “on”-state ^1^H, ^15^N assignments to the “off” state, and complemented by 3D BEST-TROSY HNcaCO, HNCACB to extend the assignment to CO, CA, and CB. Additional side chain assignments of aromatic tyrosine and histidine residues, including the tyrosine moiety of the *p*-HBI chromophore, was achieved using a series of specifically tailored 2D correlation experiments as described recently (31). Partial methyl (^1^H, ^13^C) assignments were obtained by a combination of 3D CcoNH-TOCSY, 3D HccoNH-TOCSY, and 3D H^met^C^met^C experiments, complemented by amino-acid-edited SOFAST ^1^H-^13^C correlation spectra (32, 33)

^15^N relaxation experiments (T_1_, T_2_, HETNOE) were performed using standard pulse schemes implemented in NMRlib (27). Relaxation data for 195 “on”-state and 175 “off”-state backbone amide sites could be obtained at 40 °C, from cross peaks that were well resolved and of sufficient signal to noise in the recorded 2D ^1^H–^15^N spectra, while all others (overlapping or weak) peaks were excluded from further analysis. The overall correlation time has been estimated from the “plateau” T_1_ and T_2_ values (34) as 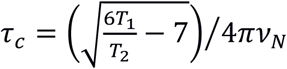.

For H/D exchange measurements, lyophilized samples of protonated ^13^C/^15^N-labelled rsFolder were dissolved in 300 μl D_2_O (sample concentration of 200 μM), transferred to a 5 mm Shigemi tube with the optical fiber inserted, and quickly put into the NMR spectrometer for data recording. The decay of amide ^1^H NMR signals, due to the exchange with deuterons, was then monitored for about 20 hours by a series of ^1^H-^15^N BEST-TROSY spectra with a time resolution of 2.5 min per experiment. The dead time of these kinetic experiments due to sample preparation, and spectrometer setup (locking, shimming, …) was about 5 min. To estimate the accuracy and reproducibility of the measurements, two data sets were recorded for both “on” and “off” states (at 600 and 850 MHz). In order to obtain more quantitative data for amide sites that are slowly exchanging in the “on” state, but much faster in the “off” state, we performed an additional H/D exchange experiment. In this experiment, the exchange measurement starts with the “on” state, and the sample is switched after a few minutes to the “off” state using high-power laser (488 nm) illumination.

## Results

### Laser-driven photoswitching

In order to be able to perform a set of time-consuming multi-dimensional (2D and 3D) NMR experiments for the metastable “off” state of rsFolder, we have adapted a setup similar to the one proposed by Mizuno et al. (2010) that allows to illuminate the sample in the NMR magnet during data collection with 488 nm or 405 nm laser light via an optical fiber introduced into the glass insert of a standard Shigemi tube (figure 2a). This illumination device is portable, can be used with any high-field NMR spectrometer (figure 2b), and can be synchronized with an NMR pulse sequence to switch the sample between the “on” and “off” states on demand.

**Figure 2.**
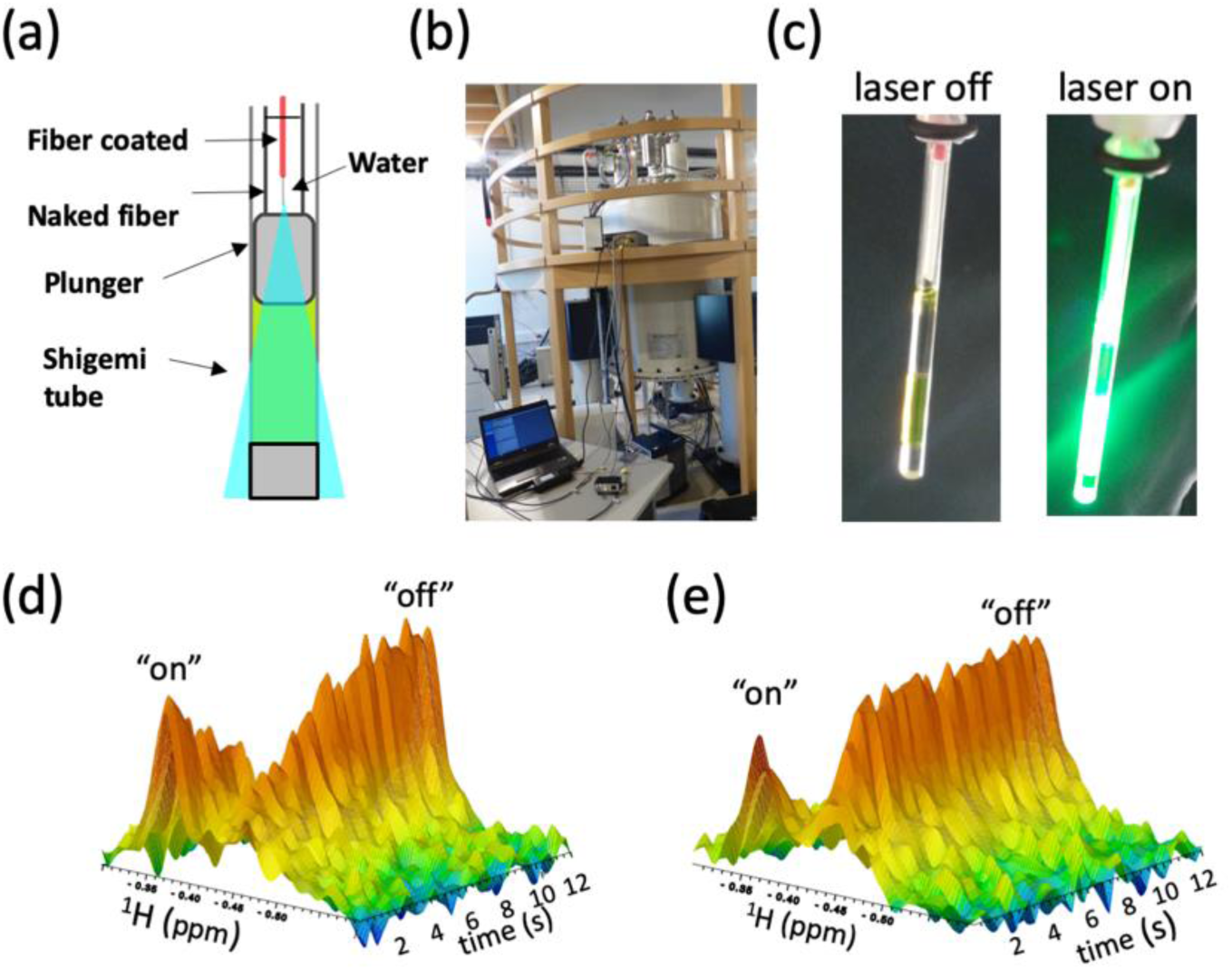
In-situ NMR sample illumination device. (a) Schematic representation of the insertion of the optical fiber into the water-filled plunger of a Shigemi NMR tube. The NMR sample is illuminated from the top in a conic geometry. (b) Photograph of the high-field NMR platform showing the portable setup of the in-situ illumination device; a laptop is connected to the laser box which is mounted on the mezzanine of the magnet. (c) Sample of rsFolder: picture with and without light illumination. (d-e) 2D plots of 1D ^1^H time series. Upon continuous laser illumination at 488 nm and power levels of (d) 70 mW and (e) 140 mW, the NMR signal of a methyl ^1^H in the “on” state decreases while the signal of the corresponding “off” state peak increases.

In order to characterize the efficiency of this illumination device to switch rsFolder between the “on” and the “off” states, we have performed real-time 1D ^1^H NMR experiments under laser illumination. The spectra plotted in Figure 2d and 2e show a decrease of the “on” state population, and an increase of the “off” state population of rsFolder over time when illuminating the sample at 488 nm. The two NMR resonances visible in this spectral window correspond to a methyl ^1^H that is spectroscopically resolved in both states, and thus presents a convenient probe for such 1D NMR-detected switching kinetics. As expected, the switching kinetics depend on the chosen laser power, with a measured on-off switching half time of about 2 s at 140 mW (maximal laser power output, Figure 2e). The lower limit of ∼2 s results from the high optical density of the sample at 488 nm, substantially increasing the time it takes for the laser light to reach all molecules in the NMR sample. Note that the inverse switching reaction, from “off” to “on” under 405 nm light illumination (120 mW) is about twice faster (data not shown).

A superposition of the ^1^H-^15^N fingerprint spectra recorded in the absence (“on” state) and presence of 488 nm light illumination (“off” state) is plotted in Figure S1. Many residues (amide groups) show significant peak shifts in these spectra, indicating that the change in chromophore geometry induced by photon absorption has a widespread effect on the amide ^1^H and ^15^N chemical shifts in the β-barrel (see below). The laser-induced peak shifts are reversible as illustrated in Figure S2 by the superposition of “on”-state and “off”-state spectra recorded before and after 1000 switching cycles using alternate sample illumination by cyan and violet light. We found that photobleaching resulted in a progressive loss of peak intensity (about 25% after 1000 cycles under the chosen experimental conditions), but did not induce any noticeable peak shifts in the spectra, indicating that the NMR-observable conformational “on” and “off” states are not affected by photobleaching. The absence of additional peaks in our NMR spectra developing over time suggests that photo-damaged rsFolder may form large aggregates not observable by liquid-state NMR.

The observed peak shifts between rsFolder “on” and “off” states offer the possibility to quantify a “switching efficiency” that can be related to the fluorescence switching contrast of importance in microscopy applications. Quantification of the residual peak intensity of “on” state peaks in the “off” state spectrum provides a measure of the relative state populations. For rsFolder, we have measured this residual “on” state population to be ∼ 3.5 % (Figure S1), independent of the sample temperature in the range 20° - 40°C. This NMR-derived value closely matches results from fluorescence spectroscopy (26), where a residual fluorescence intensity of ∼4% in the “off” state at room temperature was obtained, in line with the notion that the rsFolder “off” state is fully nonfluorescent. NMR also enables the measurement of the “off” to “on” back-switching efficiency, which in the case of rsFolder turns out be close to 100% (Figure S1).

### Laser-aided backbone resonance assignments and secondary structure

^1^H^N, 13^C^α, 13^C^β, 13^CO, and ^15^N resonance assignment of rsFolder at 40°C in the “on” state was obtained from a series of 3D BEST-TROSY HNC correlation experiments (30). Almost complete (95%) assignment was obtained for residues 2 – 230, corresponding to the β-barrel structure. Only 12 non-proline residues (W58, V69, D77, H78, E133, Y144, V151, K167, I168, K210, D211, and K215) could not be assigned from these spectra. Of those, V69, V151 and K167 are in direct contact with the chromophore based on crystallographic data. The ^1^H-^15^N correlation spectrum, annotated with the assigned residue number, is shown in Figure S3.

Recording and analyzing such a set of 3D assignment spectra is a time-consuming task (typically 1 week of data recording, and another week of data analysis). In order to speed up the assignment process for the “off” state, we have adapted a different strategy that makes use of our laser-induced in-situ switching capabilities. The “on” state assignment can be directly transferred to the “off” state by performing a laser-driven exchange experiment (LD-EXSY), conceptually similar to SCOTCH experiments (35), proposed in the past to monitor light-induced chemical reactions. The basic scheme of a LD-EXSY experiment is shown in Figure 3a. During the magnetization recovery time in the beginning of the pulse sequence (typically a few seconds) a laser illumination (488 nm) switches the proteins in the sample to the “off” state, allowing to edit either the ^1^H or ^15^N chemical shift for this state. After chemical shift editing, the magnetization is stored as longitudinal spin order (*N*_*z*_), and a second laser pulse, this time at 405 nm, converts the proteins back to the “on” state during a mixing period (exchange time) of typically a few hundred milliseconds. The remainder of the sequence allows to edit the ^1^H and ^15^N frequencies in the “on” state. The resulting 3D data set thus comprises one “off” state (^1^H or ^15^N), and two “on” state (^1^H and ^15^N) chemical shift dimensions. Figure 3b shows annotated 2D projections of two 3D data sets recorded for rsFolder, with either the ^1^H or ^15^N chemical shift labeled in the “off”-state spectral dimension. Off-diagonal peaks detected in these spectra originate from protein residues with amide chemical shifts (^1^H or ^15^N) that differ between the two states, while diagonal peaks are either due to incomplete switching, or chemical shifts that are not affected by photoswitching. These LD-EXSY spectra thus provide a straightforward and time-efficient way for amide ^1^H, ^15^N assignments in the “off” state. Additional ^13^C chemical shift information in the “off” state was then extracted from 3D BT-HNcaCO and BT-HNCACB spectra. This resulted in an assignment completeness of 93% (residues 2 – 230), with a total of 16 unassigned non-proline residues (W58, E133, Y144, S148, N150, V151, K157, Q158, N165, K167, I168, R169, L202, K210, D211, and K215), of which S148, V151 and K167 are in direct contact with the chromophore based on crystallographic data. Note that in the case of severely line-broadened amide resonances (see below) no cross peaks were observed in the triple-resonance data, and assignment was only possible with the LD-EXSY experiment.

**Figure 3.**
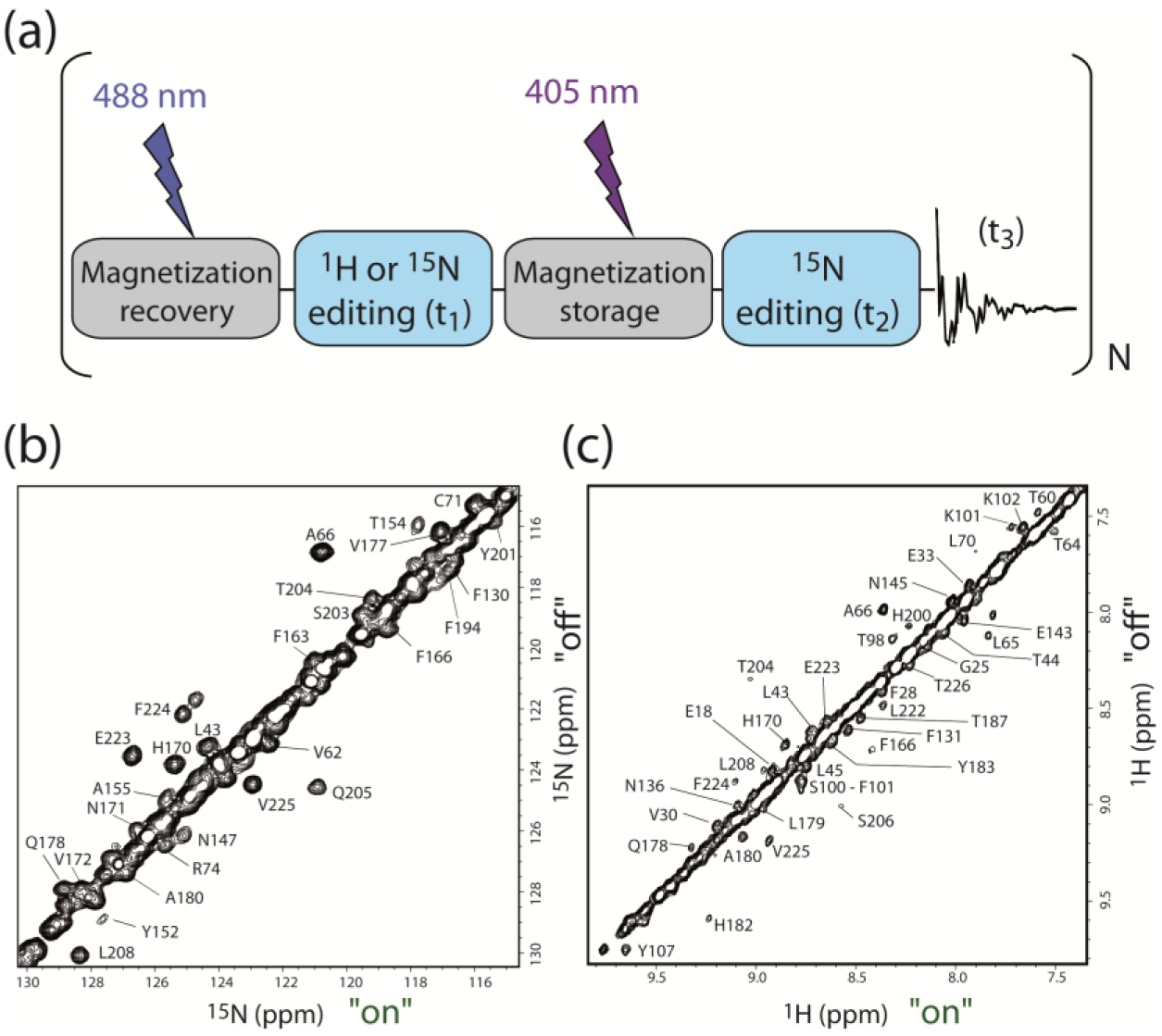
Laser-driven magnetization exchange (LD-EXSY) experiment. (a) Schematic drawing of LD-EXSY pulse sequence including light illumination at 2 different wavelengths; at 405 nm during the recycle delay (magnetization recovery, 2s), and at 488 nm during the EXSY mixing time (magnetization storage, 600 ms). The laser sources are coupled to the NMR console via TTL lines, and switched on and off by specific commands in the NMR pulse sequence code. The resulting 3D spectra have 2 “on” state and 1 “off” state chemical shift dimensions. (b) 2D projection of a 3D ^15^N-edited LD-EXSY spectrum. Off-diagonal peaks correspond to rsFolder residues (annotated) which show ^15^N chemical shift differences between “on” and “off” states. (c) Same as (b) but for ^1^H-edited LD-EXSY.

The assigned NMR frequencies, and in particular the ^13^C chemical shifts, provide useful information on the secondary structural elements (α-helix, β-strand) present in rsFolder under our experimental conditions. In Figure S4, the secondary structural features derived from our measured chemical shifts using the TALOS-N program are plotted (36). A comparison with the X-ray structures of rsFolder (PDB files 5DTZ and 5DU0) shows that the β-barrel structure in solution is almost identical to that observed in the crystal at cryogenic temperature. Nevertheless, a few slight differences are observed for strand β7 and the two helices (H2 and H3) inside the barrel to which the chromophore is connected. The β7 strand in solution is extended by 3 residues at the N-terminal end, helix H2 is shifted by 3 residues towards the C-terminus, and helix H3 has a non-canonical backbone geometry that is not identified as an α-helix by TALOS. Interestingly, it has been concluded from the hydrogen bonding patterns observed in the high resolution crystal structure of EGFP (37), an “ancestor” protein of rsFolder, that helix H2 is composed of a 3_10_ helix (P57 – L61 in rsFolder) followed by a short regular α-helix (L61-L65 in rsFolder), in agreement with our NMR data.

### Backbone dynamics in the “on” and “off” states

In order to obtain information on the local backbone dynamics we have performed ^15^N relaxation measurements of rsFolder in both the “on” and “off” states at 40°C. The measured T_1_, T_2_, and HETNOE data are plotted in Figure 4a as a function of the protein sequence, with the secondary structural elements as identified from TALOS-N plotted on top of each graph. These data indicate an overall rigid monomeric globular protein, with a tumbling correlation time of about 10 ns at 40°C. Increased flexibility on the ps to ns time scale (longer T_2_ and reduced HETNOE values) is detected in some loop regions, especially the loops connecting the first helix (H1) and strand β1, and the loop between β-strands 9 and 10. Most interestingly, there is almost no significant difference in the fast time scale dynamics, as seen from these ^15^N relaxation data, between the “on” and “off” states. However, some differences are observed that can be ascribed to increased conformational dynamics on the µs to ms time scale in the “off” state of rsFolder. A first observation is the lack of peak intensity for some amides in β-strands 7 and 8 in the ^15^N relaxation spectra (turquoise bars in Figure 4b), as well as reduced T_2_ values measured for residues C71 and S73 in the non-canonical helix H3. The reduction in peak intensities for some residues in the “off” state can also be observed in standard 2D ^1^H-^15^N BEST-TROSY spectra, as illustrated in Figure 4b. In order to further characterize these conformational dynamics we have performed CPMG-type relaxation-dispersion experiments. Unfortunately, the signal to noise ratio obtained for the above-mentioned residues was too low to perform a quantitative analysis. Only S73 yielded data of reasonable quality (Figure 4c) that confirm the presence of a conformational exchange process in the “off” state that leads to a modulation of the chemical shift during the relaxation period, while no relaxation dispersion is observed for this residue in the “on” state. Due to the limited experimental sensitivity, we did not attempt to extract kinetic (K_ex_) or thermodynamic (state populations) information from these dispersion curves. As the exchange contribution to the transverse relaxation rate is not completely quenched at the highest CPMG frequency of 1000 Hz, we can reasonably assume that the exchange process is faster than 1000 s^-1^.

**Figure 4.**
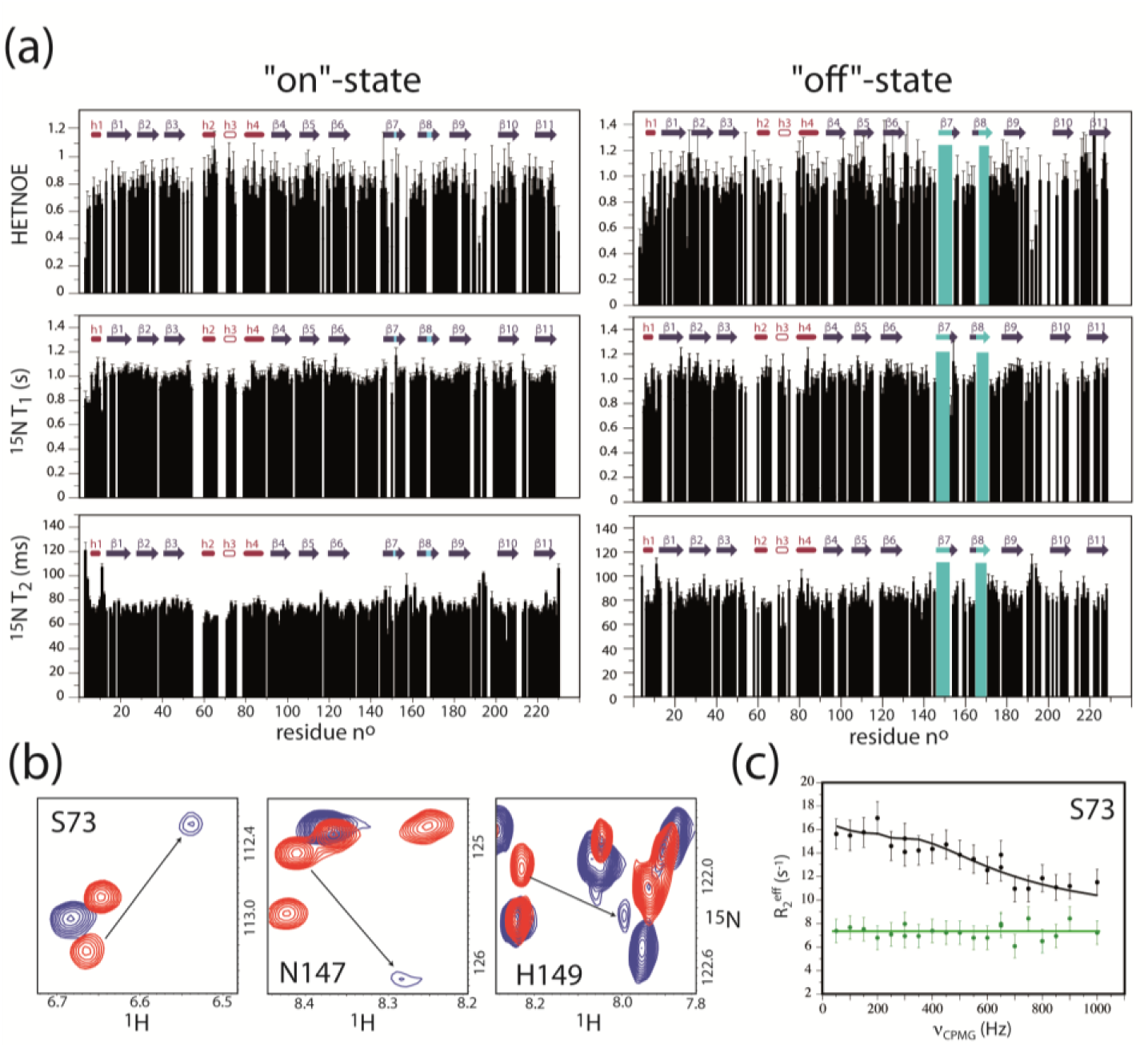
Backbone dynamics of rsFolder in “on” and “off” states. (a) ^15^N relaxation data (T_1_, T_2_ and HETNOE) measured at 700 MHz and 40°C, plotted as a function of protein sequence. Secondary structural elements as identified from the ^13^C chemical shifts by the program TALOS-N are plotted on top of each graph. Cyan bars indicate peptide region for which no peak is detected in the “off” state. (b) ^1^H-^15^N BEST-TROSY spectral regions (red: “on” state, blue: “off” state) zooming on line-broadened “off” state peaks. (c) CPMG-RD data recorded for residue S73 at 850 MHz in the “on” (green) and “off” (black) states.

Figure 5 summarizes the main changes in NMR observables after light illumination. The following NMR data have been color-coded on the structure of rsFolder in an open-sheet and a 3-dimensional representation: (i) unobserved amide resonances in both the “on” and “off” states (dark blue); (ii) amide resonances undetected in the “off” state only (blue); (iii) amide resonances with significantly reduced peak intensity in the “off” state as extracted from the graph of figure S5b, and/or decreased ^15^N T_2_ values (light blue); (iv) significant ^1^H-^15^N chemical shift changes (Δ_CS_ > 2 ppm) as extracted from the graph of figure S5a (orange). Note that the reported chemical shift changes are not a direct evidence of changes in structure or conformational dynamics, as part of it may be due to long-range ring-current-induced chemical shift effects that follow from the change in orientation of the chromophore. Interestingly, all of these NMR data point towards significant changes in conformational dynamics in a particular β-barrel region. The missing “on” state peaks for a few residues in β7 (V151) and β 8 (K167, I168) indicates some possible dynamics already present in the fluorescent state, highlighting a well-known defect in the β-barrel structure and a hotspot for backbone fluctuations (16, 38, 39). After the *cis-trans* isomerization of the chromophore, the conformational dynamics occurring on a μs to ms time scale are expanded to a larger region close to the phenol ring of the chromophore, comprising strands 7, 8, 10, and eventually 11, as well as helix H3.

**Figure 5.**
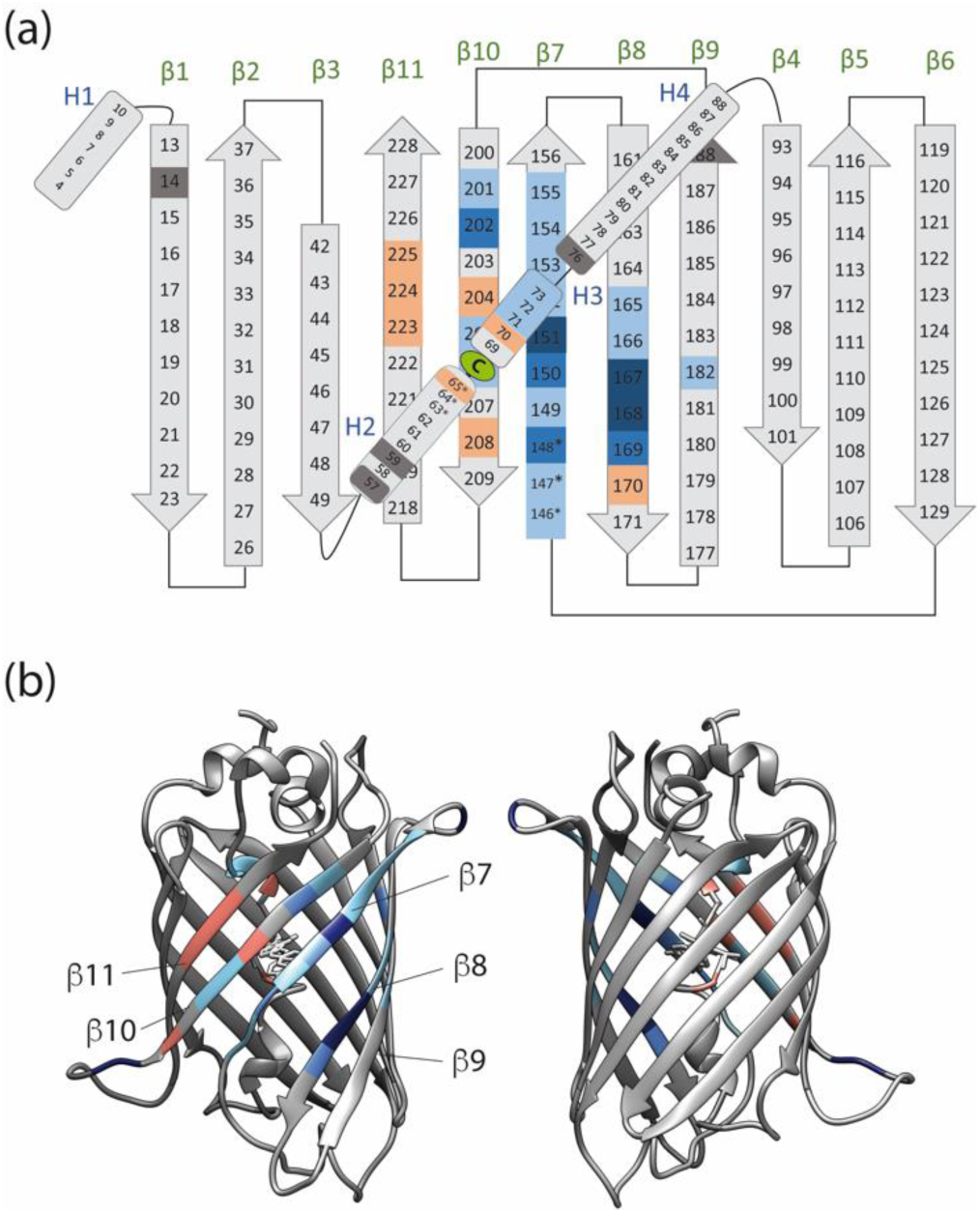
Increased conformational dynamics on the μs-ms timescale in the “off” state of rsFolder. The NMR-derived information is color-coded on the (a) open-sheet and (b) 3-dimensional representation of the rsFolder structure as revealed by X-ray crystallopgraphy and NMR (A ‘*’ indicates structural elements only identified by NMR): (i) undetected/unassigned amides in both states (dark blue), (ii) undetected/unassigned amide peaks in the “off” state only (middle blue), (iii) strongly line-broadened amide NMR signals in the “off” state (light blue), (iv) amide 1H-15N with large chemical shift changes between “on” and “off” states (orange). In addition, proline residues lacking an amide proton are color coded in dark grey.

### Fold stability and water accessibility

Complementary information on the conformational dynamics of backbone amides in the “on” and “off” states of rsFolder has been obtained from hydrogen-deuterium (H/D) exchange measurements. For amide hydrogens engaged in an intramolecular H-bond, H/D exchange is assumed to proceed via transient opening of the hydrogen bond followed by interaction with a solvent exchange catalyst (40). Therefore, NMR derived H/D exchange rates provide an indirect measure of the dynamic fluctuations of the protein backbone leading to transient breakage of hydrogen bonds. At neutral or acidic pH, the formation of the H-bond after transient opening is generally much faster than the intrinsic chemical exchange process. In this so-called EX2 exchange regime, the exchange rate constants provide a measure of the exchange-competent state population. A second structural parameter that needs to be considered is water accessibility. Typically, an amide hydrogen surrounded by water has a higher exchange probability than the same amide hydrogen buried in the interior of a globular protein. For rsFolder, the available crystal structures for the “on” and “off” states do not indicate any major change in water-accessible cavities inside the β-barrel as illustrated in Figure S7. Therefore we consider that changes in the observed H/D exchange rates are mainly due to a population increase (or decrease) of exchange competent, unprotected conformational sub-states.

H/D exchange kinetics for hydrogen-bonded amides of rsFolder were measured by time-resolved 2D ^1^H-^15^N NMR. The exchange process was initiated by dissolving the lyophilized protein in D_2_O outside the NMR instrument. Typically, unprotected solvent-accessible amide sites were fully exchanged during the experimental dead time of about 5 minutes. The exchange process was then monitored for about 20 hours, allowing to quantify exchange processes with time constants T_ex_=1/k_ex_ spanning a range of 3 orders of magnitude (5 min < T_ex_ < 5000 min). The measured exchange time constants are color-coded (logarithmic scale) on the secondary structure (open sheet representation) of rsFolder in Figure S6 for the “on” and “off” states. Examples of intensity decays extracted from the real-time NMR data for individual amide sites in the “on” and “off” states are shown in Figure 6a. A first observation from these H/D exchange data is that, while overall the amide protons in the β-barrel of the “on” state are well protected from solvent exchange, strand β7 is much less protected due to the lack in hydrogen bonding with the adjacent β8 strand. In the “off” state, a significant increase in the measured solvent exchange rates is detected for amides in the neighboring strands β8 and β10, further emphasizing the increased dynamics in this part of the β-barrel structure upon light illumination, leading to a destabilization of the hydrogen bonding network. Some local destabilization is also observed for amide hydrogens in strands β4 and β5, while no significant changes are observed for the remainder of the β-barrel (β1-3, β6, β11) as illustrated in Figure 6b. The most pronounced changes in protection from solvent exchange, however, are observed for the two helices (H2 and H3) flanking the chromophore in the interior of the β-barrel (see Figure 6b), with a decrease in solvent protection ranging from a factor of 5 up to a factor of 100. These data clearly indicate the presence of conformational dynamics affecting these helical elements induced by the chromophore *cis-trans* isomerization. Interestingly, while ^15^N relaxation and line broadening data already identified increased dynamics in helix H3, they did not point towards changes in helix H2 dynamics. Our findings highlight the complementarity of NMR relaxation and H/D exchange data in providing information on local protein dynamics, the latter being sensitive to much longer time scales and smaller chemical shift changes.

**Figure 6.**
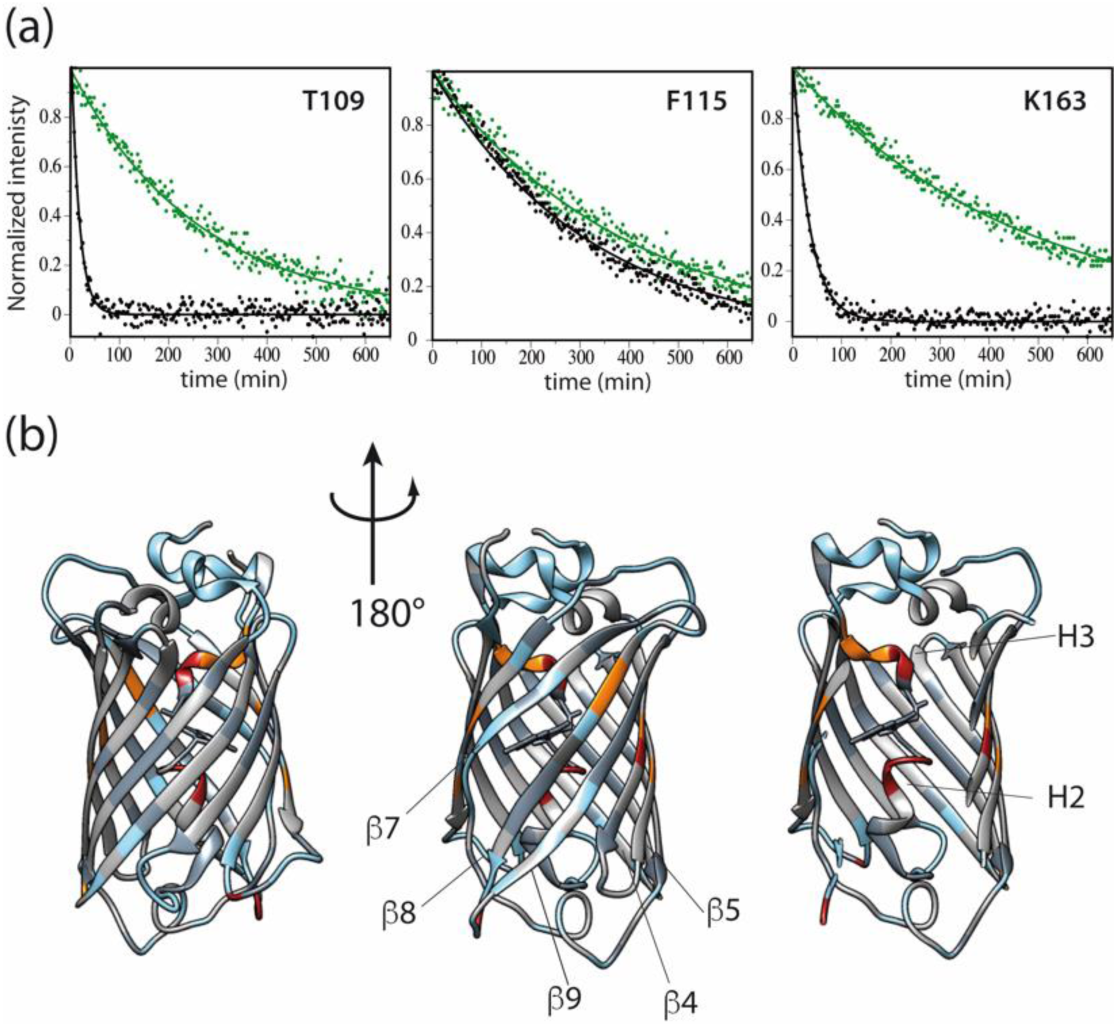
H/D exchange NMR data. (a) Representative intensity decay for individual amide hydrogens extracted from real-time 2D NMR data recorded for the “on” (green) and “off” (black) states of rsFolder at 40°C and 850 MHz ^1^H frequency. (b) The derived H/D exchange information is color-coded on the crystal structure of rsFolder: (i) residues with amide hydrogen exchange time constants Tex < 5 min in both the “on” and off” states (light blue); (ii) residues for which the hydrogen exchange rate is increased by a factor of 4-20 (orange); (iii) residues for which the hydrogen exchange rate is increased by more than a factor of 20 (red); (iv) residues without a significant change in exchange rate between “on” and “off” states (grey); (v) no information due to peak overlap, proline, or unassigned residues in one of the two states (dark grey).

### Sidechain and chromophore dynamics in the “on” and “off” states

Line broadening effects similar to those already discussed in the context of backbone dynamics (Figure 4b) are also observed in the ^1^H-^13^C correlation spectra of rsFolder in the “off” state, indicating that a number of protein side chains undergo conformational exchange. Examples of methyl groups (I168 and T63) and an aromatic side chain (H149) that become severely line broadened in the “off” state are shown in Figure 7a. In order to identify the protein regions that are involved in conformational side chain dynamics, we have performed ^1^H, ^13^C resonance assignment of methyl and aromatic side chain moieties in the “on” and “off” states of rsFolder as further explained in the Materials and Methods section. Figure S8 highlights ^1^H-^13^C spectral regions displaying NMR signals that experience either large chemical shift changes or severe line broadening in the “off” state. The side chains involved in conformational dynamics are all located in close proximity to the chromophore (Figure 7b), attached to β-strands 7, 8, 10, and 11 or helices 2 and 3, the same backbone regions that have been shown to experience conformational exchange in the “off”-state. These data further support the conclusion that the chromophore itself and/or the surrounding environment experience significant conformational dynamics in the “off” state leading to chemical shift modulation on the μs to ms time scale.

**Figure 7.**
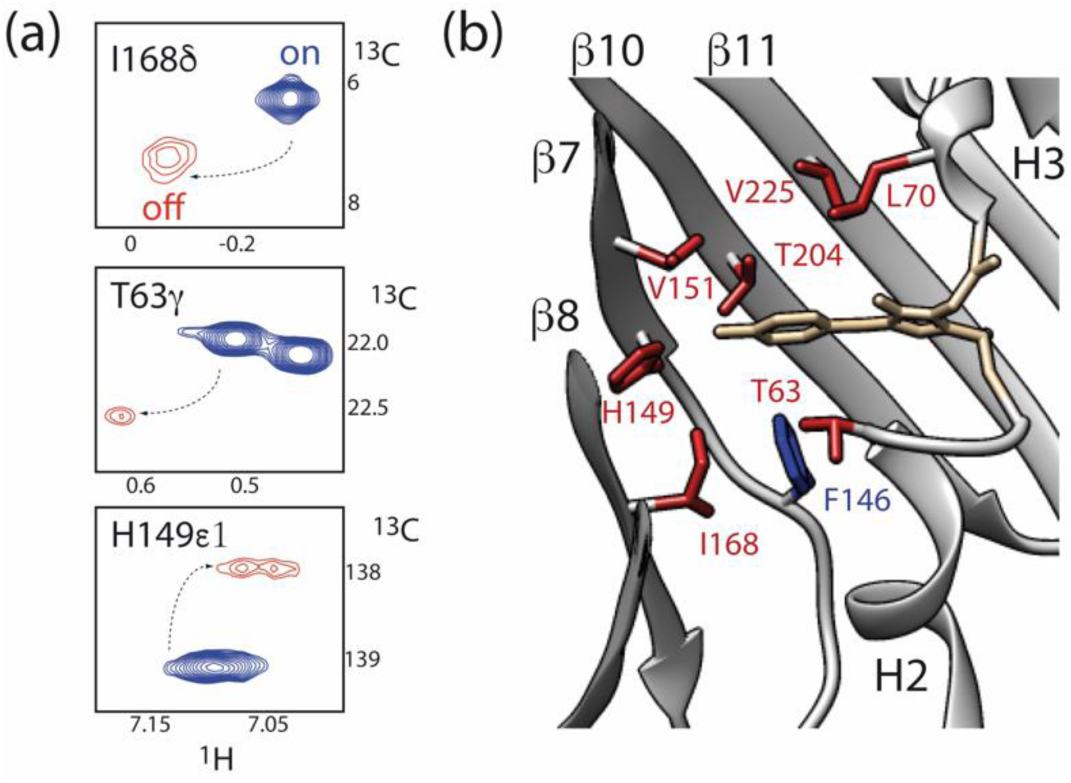
rsFolder side chains involved in conformational exchange processes in the “off” state as deduced from extensive line broadening. Examples of peak line broadening observed in the ^1^H-^13^C correlation spectra of rsFolder: on-state (blue), “off”-state (red). (b) Zoom on the chromophore environment highlighting the side chains which show NMR line broadening in the “off” state (red). In addition, F146 is plotted as it experiences large chemical shift changes upon light illumination.

NMR also provides information on the dynamics of the chromophore, and its stabilization in the β-barrel by hydrogen bonding. The exceptional thermal stability of rsFolder in the “off” state was tentatively attributed to the H-bond that H149 maintains with the phenol ring of the chromophore in both ‘on’ and ‘off’ states (26), as illustrated in Figure 8a. In order to validate or contradict this hypothesis, we first performed NMR experiments that reveal the protonation state of H149. In the “on” state, the imidazole ring of H149 is predominantly in a neutral tautomeric state, protonated at the Nδ_1_ position (31). In the “off” state, as discussed in the previous paragraph, the ^1^H-^13^C correlation peaks of H149 (Hδ_1_-Cδ_1_ and Hε_2_-Cε_2_) are severely line broadened, and no correlation peaks are detected in aromatic HCC-type experiments. Therefore, no conclusions can be drawn on the protonation state of H149 in the “off” state. Further information on the involvement of H149 in hydrogen bonding interactions was obtained by a ^1^H-^15^N SOFAST-HMQC experiment (32) tailored to the chemical shifts of histidine side chains. The presence of a hydrogen bond protects the “labile” imidazole hydrogen(s) from solvent exchange. Therefore, a correlation peak for the protonated histidine ring nitrogens is expected if the lifetime of the hydrogen is at least a few hundred milliseconds. Figure 8b shows such correlation spectra recorded for the “on” and “off” states at different temperatures (15°C, 25°C, and 40°C). At low temperature (15°C), a total of 6 out of the 10 histidines of rsFolder give rise to a cross peak in the “on” state indicating their involvement in a hydrogen bond. While 4 of them (H26, H182, H200, and H218) remain visible also at higher temperature (40°C), no correlation peak is detected for H149 and H232 at this temperature. Taking H218 as an internal reference, the relative H-bond stability (peak intensity) decreases by a factor of 2 when increasing the temperature from 15° to 25°C. Our NMR data validate the crystal-structure-derived hypothesis of H-bond formation between the Nδ_1_ of H149 and the phenol ring of the chromophore. However, we can also conclude that this H-bond is relatively unstable, as compared to other H-bonds involving histidines H26, H182, H200, and H218. In the “off” state, the same peak patterns and T-dependence is observed, except for H149 that does not give rise to a correlation peak at any temperature. This observation, however, does not allow to conclude on the absence of a H-bond between H149 and the chromophore in the “off” state, as the protonated phenol ring has to interact with an unprotonated imidazole ring nitrogen in order to form an H-bond, that is undetectable in our NMR spectra. Nevertheless, the observed line broadening of the non-exchangeable ring hydrogens of H149, resulting from conformational exchange dynamics, makes the hypothesis of a stable H-bond in the “off” state very unlikely.

**Figure 8.**
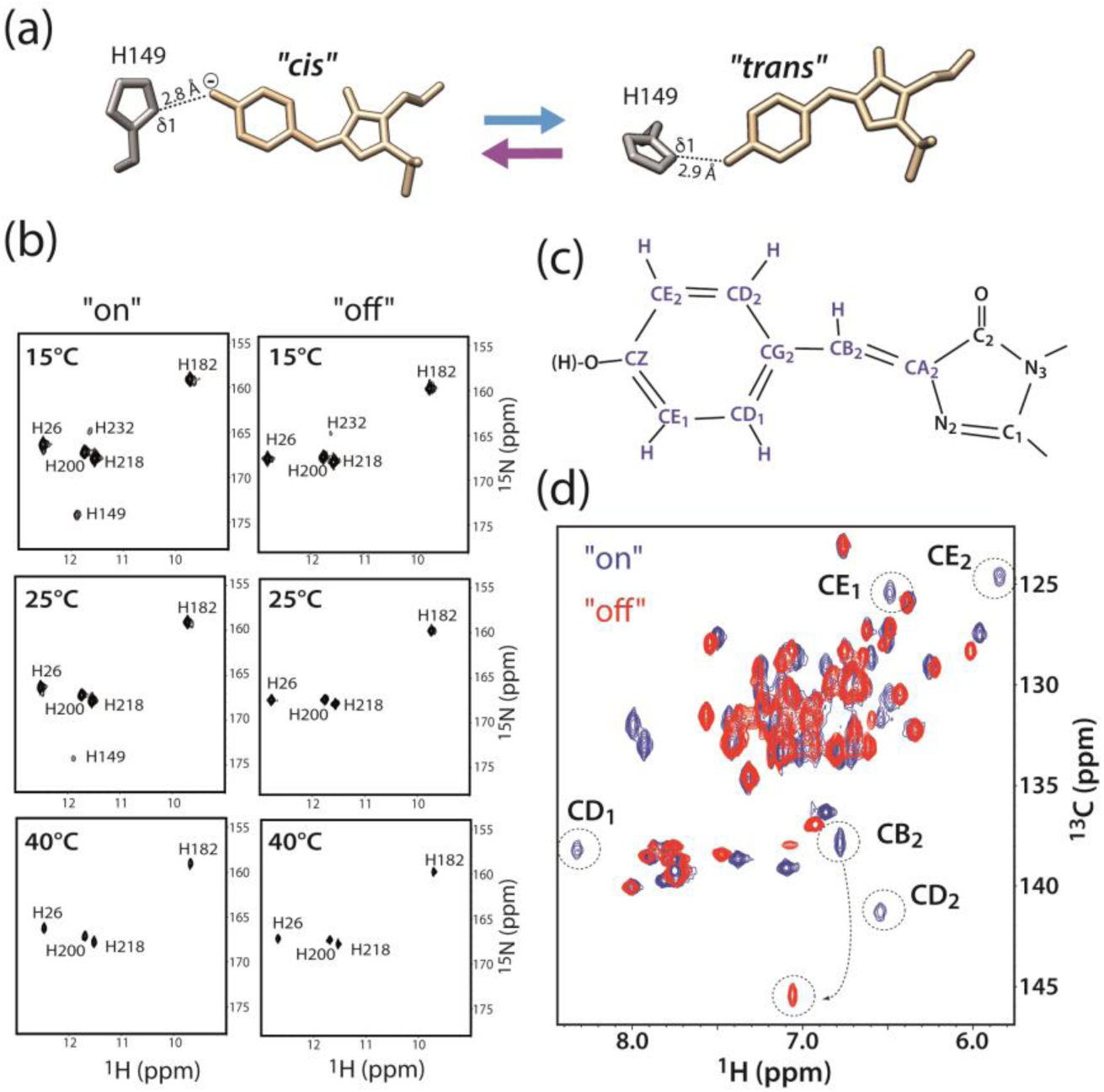
NMR information of the chromophore. (a) Chromophore and His149 side chain conformations extracted from the available crystal structures of rsFolder in the “on” (*cis* configuration) and “off” (*trans* configuration) states. (b) Temperature series of ^1^H-^15^N SOFAST-HMQC spectra of histidine side chains in the “on” and the “off” states recorded at 850 MHz ^1^H frequency. The detected cross peaks are annotated by the residue number. (c) Schematic representation of the chemical structure of the chromophore. Nuclei giving rise to NMR signals are highlighted by blue letters, and their NMR frequencies are provided in Table 1. (d) Overlay of 2D aromatic ^1^H-^13^C Best-HSQC spectrum (31) of rsFolder, recorded without CT ^13^C editing to achieve highest experimental sensitivity, in the “on” (blue) and “off” (red) state. NMR signals from the tyrosine moiety (Y63) of the chromophore are annotated.

We also attempted to obtain some information on the NMR signature of the chromophore itself. Figure 8d shows the aromatic region of the ^1^H-^13^C correlation spectra of rsFolder in the “on” state (blue spectrum). The chromophore contains 5 C-H pairs (Figure 8c) in the tyrosine (Y) part of the chromophore that is made up from the A66-Y67-G68 tripeptide in the rsFolder sequence. The 5 corresponding correlation peaks detected in the spectrum of Figure 8d were identified and unambiguously assigned by a set of 2D correlation spectra shown in Figure S9, recorded with pulse sequences that were specifically tailored for tyrosine side chains (31). These NMR data allowed us to assign all ^1^H and ^13^C resonances (except for CO) of the tyrosine moiety of the chromophore (Table 1). The upfield shifted CZ chemical shift (178 ppm) is characteristic of an anionic phenol group as expected for a deprotonated chromophore in the fluorescent “on” state. The observation of distinct correlation peaks for the symmetric CD and CE sites of the phenol ring indicates that the chromophore is structurally stabilized in the β-barrel with slow ring flip dynamics (k_ex_ < 500 s^-1^) at 40°C. These slow ring flips are in contrast to all other tyrosine and phenylalanine side chains of rsFolder for which a single correlation peak was detected for the symmetric δ1 and δ2 (ε1 and ε2) sites, indicative of fast ring flip dynamics leading to chemical shift averaging. In the non-fluorescent “off” state (red spectrum in Figure 8d), the 4 correlation peaks of the chromophore’s phenol group disappear, indicative of a significant change in chromophore dynamics. Only the signal of the HB-CB in the methine bridge of the chromophore remains detectable in the dark state, with the CB resonance experiencing an impressive 7.6 ppm upfield shift after *cis-trans* isomerization of the chromophore. The NMR assignment of the chromophore in the “off” state could be further extended to the CA and CG (Figure S9b) that also shift upfield by 3.3 and 4.4 ppm, respectively (see table 1). Our NMR data indicate a destabilization of the chromophore in the “off” state leading to motion of the phenol ring on the millisecond time scale. However, whether these motions correspond to increased ring flip dynamics, local fluctuations around the methine bridge, or both cannot be distinguished from these data.

**Table 1.**
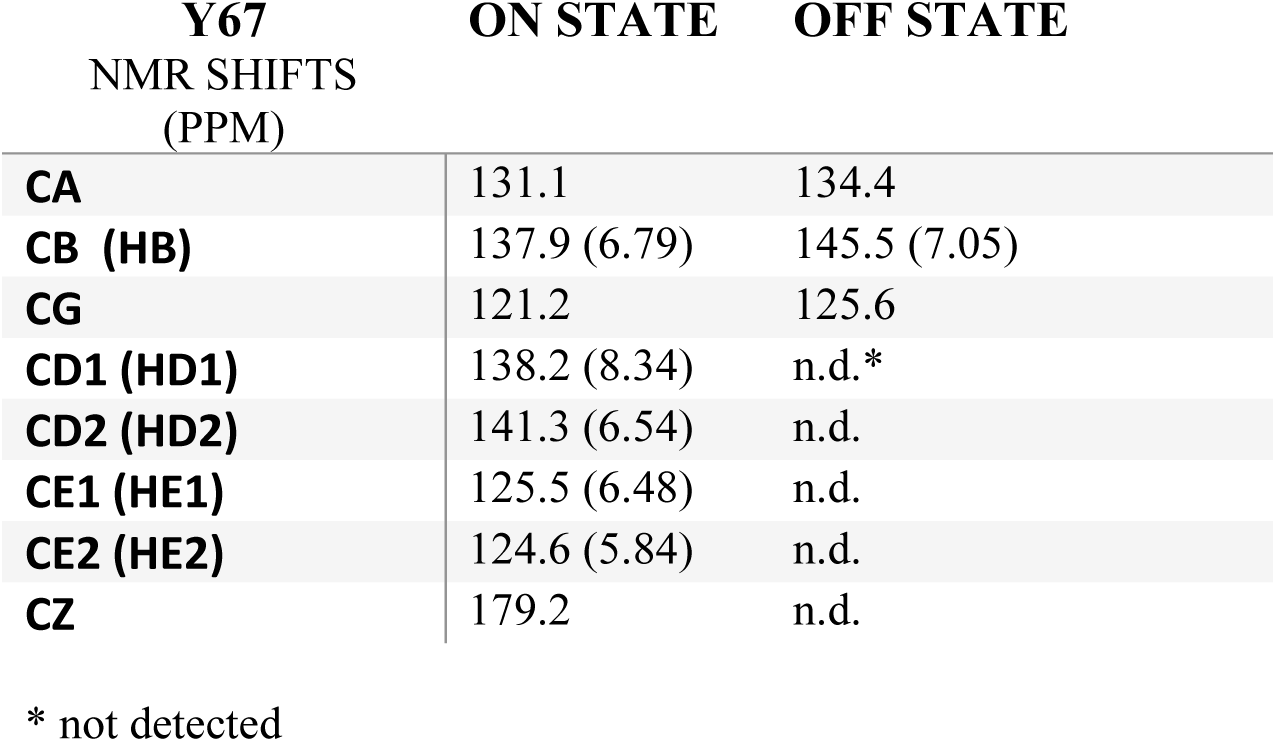
Chromophore ^1^H and ^13^C chemical shifts

## Discussion

### In-situ NMR illumination and time-efficient NMR experiments

Our portable laser illumination device allowed us to perform a large set of 2D and 3D correlation experiments on a single rsFolder sample. Due to its portability, the setup is compatible with any high-field NMR instrument and probe, provided that the optical fiber is long enough for the sample to reach its position in the magnet. Illumination from the top (fiber placed in the Shigemi plunger) ensures minimal perturbation of the magnetic field homogeneity, and thus results in close to optimal NMR line shapes. However, relatively high laser power is required to ensure fast “on” to “off” (or “off” to “on”) switching, in particular due to nonuniform illumination and high optical density of the sample. The LD-EXSY experiment that makes use of fast switching capabilities provides a convenient and time-efficient way to transfer NMR assignments of fast-switching PTFPs from one conformational state to another. This is of importance as NMR resonance assignment is the most time-consuming step for investigating the dynamic properties of PTFPs. Once a sufficient level of resonance assignment has been achieved, NMR provides a unique tool to access local dynamics in the protein backbone, side chains, and chromophore on a broad range of time scales, as will be discussed further in the following paragraph(s).

### Conformational dynamics in the fluorescent state of rsFolder

The β-barrel scaffold of fluorescent proteins stabilizes the anionic *cis* conformation of the chromophore via hydrogen bonding, π-stacking and cation-π interactions, as well as steric hindrance. This restricts the chromophore’s internal motion observed in free solution, and prevents radiationless decay upon light excitation, a prerequisite for a high fluorescence quantum yield. However, this conformational rigidity is somehow in contradiction with the capability of RSFPs to switch efficiently between distinct conformations, implying local structural rearrangements, as well as changes in the hydrogen bonding network upon light excitation. Therefore, a detailed experimental characterization of the structural heterogeneity and conformational flexibility of the chromophore and its protein environment is important for understanding and improving the photochromic properties of RSFPs. Such dynamic information is typically not available from crystallographic structures. In the case of rsFolder, the available X-ray structures show a well-defined β-barrel fold, and a single chromophore conformation in both the “on” and “off”-states. Here, we have applied liquid-state NMR spectroscopy to complement this static picture. Information on the local dynamics over time scales ranging from picoseconds to about a few hundred milliseconds have been obtained from a combination of ^15^N spin relaxation measurements, NMR line width analysis, and H/D exchange data. Our NMR data point towards a well-known defect in the β-barrel structure of GFP-derived fluorescent proteins around strand β7 (16, 18, 38, 41) where the H-bonds formed with the neighboring strands (especially β8), typical of an anti-parallel β-sheet conformation, are weakened by longer distances and other structural irregularities. This leads to backbone fluctuations and transient opening of the β-barrel in the fluorescent “on” state of rsFolder as evidenced by a few unobservable amide resonances (V151, K167, and I168), and fast H/D exchange rates measured for amide hydrogens in strand β7. Similar observations have been made previously for conventional GFPs (16, 18). The dynamic “holes” in the β-barrel between β7 and the neighboring strands have also been identified by MD simulations as major entry points of water molecules into the interior of the β-barrel (39). Although certainly not a sufficient condition, such conformational plasticity is likely required for efficient photoswitching.

The data show clear evidence of an anionic chromophore, with the phenolate oxygen hydrogen-bonded to the Nδ1 of H149. This finding is in agreement with the crystal structure (26) that indicates that the chromophore’s phenolate oxygen is involved in 3 hydrogen-bond interactions with H149, T204, and a water molecule. However, our NMR data show that the H-bond between the chromophore and H149 is thermally unstable. At physiological temperature, the bond is constantly broken and reformed on a sub-millisecond time scale. It has been shown by QM/MM simulations of the RSFP Dronpa (42) that two or three H-bonds stabilizing the chromophore are required for high fluorescence quantum yields, while less hydrogen bonding favors photo-switching of the chromophore by a hula-twist mechanism. We speculate that the weak H-bond observed with H149 in rsFolder is a characteristic property of RSFPs that have been engineered to obtain reasonable fluorescence quantum yields combined with efficient switching capability.

### Light-induced conformational dynamics in the chromophore and its environment

As for a majority of RSFPs, on-off photoswitching in rsFolder involves a *cis*-*trans* isomerization of the chromophore coupled to deprotonation at the hydroxy group of the phenol moiety. In our NMR spectra, this photoinduced isomerization results in large upfield ^13^C shifts of carbons in the methine bridge (CA, CB, and CG) of the chromophore. As ^13^C chemical shifts are particularly sensitive to the local geometry (torsion angles) of the chemical bonds (43), the methine bridge ^13^C chemical shifts can be considered a spectral signature of *p*-HBI being in *cis* or *trans* configuration. In principle, the protonation state at the chromophore’s phenol moiety can be inferred from the ^13^C_Z_ chemical shift, which is expected to be 175-180 ppm for a phenolate, and 155-160 ppm for a phenol. However, for rsFolder in the “off” state no NMR signals could be detected for the phenol moiety in the aromatic ^1^H-^13^C spectra, indicative of a chromophore undergoing conformational exchange dynamics on the μs to ms time scale. Similarly, NMR signals of nuclei in the protein side chains H149 and T204, forming hydrogen bonds with the phenolate oxygen in the “on” state”, become strongly line broadened in the “off” state. These NMR observations clearly indicate a destabilization of the chromophore, and the absence of any stable hydrogen bonding interaction with the β-barrel. In the crystal structures of rsFolder, the observed distances between the histidine δ1 nitrogen and the phenol oxygen are only little altered (from 2.8 Å to 2.9 Å) upon photo-isomerization of the chromophore (Figure 7). Our findings thus contradict the conclusion drawn from these structures that H149 remains hydrogen bonded to the hydroxyl group of the phenol in the “off” state. Most likely, the crystal structure observed at cryogenic temperatures represents the lowest energy conformation in the “off” state, which at higher temperature is in exchange with alternate conformations that are separated by low energy barriers. A loss of hydrogen bonding of the chromophore phenolate with the β-barrel scaffold has also been observed for other RSFPs, e. g. Dronpa, rsEGFP2, and rsGreen (8, 42, 44, 45). Of note, this has also been observed recently in the case of a long-lived dark state in the photoconvertible fluorescent protein mEos4b (7).

The chromophore cis-trans isomerization also induces increased structural dynamics at the β-barrel side (β7 – β10) facing the chromophore’s phenol group, the two helical regions (H2 and H3) connecting the chromophore with the rest of the protein, and a number of side chains attached to these peptide regions, and pointing towards the chromophore. In particular, all side chains that undergo a structural rearrangement in the “off” state (T63, F146, H149, V151, and I168), as seen by X-ray crystallography (Figure 1), are heavily line-broadened in our NMR data (Figure 6) except for F146 that shows large chemical shift changes. These observations further consolidate the hypothesis that photo-isomerization in rsFolder leads to a polymorphic and highly dynamic chromophore environment with multiple sub-states (conformations) of comparable free energy that interconvert on the micro-to-millisecond time scale. Most likely, the structural fluctuations of the chromophore and surrounding side chains also effects the backbone regions to which they are attached, thus explaining the observed amide ^1^H line broadening and increased solvent exchange rates. These data are fully in line with the notion of a fully nonfluorescent dark state unable to undergo radiative de-excitation upon photon absorption. Similar findings on changes in backbone dynamics upon photo-switching have previously been reported for the anthozoan-derived RSFP Dronpa (14).

So far, the chromophore connecting helical elements, H2 and H3, have found only little attention in the field of GFP engineering. Concerning the helical region H3, we know that L70 is an important residue for the photo-switching properties of rsEGFP2 (24) and rsFolder (26), while S72 (S73 in rsFolder) has been suggested as one of the ‘hopping’ points of protons when entering from the solvent onto their way to the chromophore (39). In our NMR data, the H3 region of rsFolder shows increased conformational dynamics in the “off” state, as evidenced by line broadening (C71, F72, S73) and faster H/D exchange (L70, C71). It has also been reported recently using time-resolved serial crystallography at an XFEL that during photo-switching in rsEGFP2 an intermediate state becomes populated, in which the chromophore is twisted, and helix H2 is moved downward along the β-barrel axis (8). Here, we have shown that water exchange for several amide hydrogens (V62, T63, and T64) in H2 is increased by up to 2 orders of magnitude. We may therefore speculate that conformations similar to those seen in the excited state may be also transiently occupied under equilibrium conditions at ambient temperature, possibly involved in the thermally induced relaxation mechanism of rsFolder from the “off” state to the “on” state.

## Conclusions

In this work, we have used multidimensional liquid-state NMR spectroscopy coupled to in-situ sample illumination as a complementary high-resolution tool to study light-induced changes in the conformational dynamics of the photo-transformable fluorescent protein rsFolder. Our results add a dynamic dimension to the static view provided by X-ray crystallographic protein structures. In particular, we identified dynamic hotspots in the chromophore environment and the β-barrel structure that may suggest future mutation sites to be probed by e.g. saturation mutagenesis in order to alter the photophysical properties of RSFPs. Furthermore, NMR spectroscopy provides a unique tool to investigate the influence of environmental conditions on the conformational dynamics of PTFPs, and correlate them with their photophysical and photochemical properties. Therefore, including NMR spectroscopy into the toolbox used for rational fluorescent protein engineering will provide new opportunities for further improving advanced fluorescent markers for a wide range of microscopy and biotechnological applications.

## Author Contributions

DB and BB conceived and directed the research; IA and KG expressed the protein and prepared the samples; NC and BB performed the NMR experiments and analyzed the data; VA and MB contributed plasmids, and helped with the experimental setup; NC, DB and BB wrote the paper. All authors reviewed the results and approved the final version of the manuscript.

## Acknowledgements

We thank Dr Paul Schanda for many stimulating discussions. Financial support from the CNRS (Défis Instrumentation 2018) is acknowledged. This work used the NMR and isotope labeling platforms of the Grenoble Instruct-ERIC center (ISBG; UMS 3518 CNRS-CEA-UJF-EMBL) within the Grenoble Partnership for Structural Biology (PSB). Platform access was supported by FRISBI (ANR-10-INBS-05-02) and GRAL, a project of the University Grenoble Alpes graduate school (Ecoles Universitaires de Recherche) CBH-EUR-GS (ANR-17-EURE-0003). IBS acknowledges integration into the Interdisciplinary Research Institute of Grenoble (IRIG, CEA).

